# A prostate cancer gastrointestinal transcriptional phenotype may be associated with diminished response to AR-targeted therapy

**DOI:** 10.1101/2024.06.02.595931

**Authors:** Aishwarya Subramanian, Meng Zhang, Marina Sharifi, Thaidy Moreno-Rodriguez, Eric Feng, Nicholas R. Rydzewski, Raunak Shrestha, Xiaolin Zhu, Shuang G. Zhao, Rahul Aggarwal, Eric J. Small, Chien-Kuang Cornelia Ding, David A. Quigley, Martin Sjöström

## Abstract

**Background:** Prostate cancer is a heterogenous disease, but once it becomes metastatic it eventually becomes treatment resistant. One mechanism of resistance to AR-targeting therapy is lineage plasticity, where the tumor undergoes a transformation to an AR-indifferent phenotype, most studied in the context of neuroendocrine prostate cancer (NEPC). However, activation of additional de- or trans-differentiation programs, including a gastrointestinal (GI) gene expression program, has been suggested as an alternative method of resistance. In this study, we explored the previously identified GI prostate cancer phenotype (PCa-GI) in a large cohort of metastatic castration-resistant prostate cancer (mCRPC) patient biopsy samples.

**Methods:** We analyzed a dataset of 634 mCRPC samples with batch effect corrected gene expression data from the West Coast Dream Team (WCDT), the East Coast Dream Team (ECDT), the Fred Hutchinson Cancer Research Center (FHCRC) and the Weill Cornell Medical center (WCM). Survival data was available from the WCDT and ECDT cohorts. We calculated a gene expression GI score using the sum of z-scores of genes from a published set of PCa-GI-defining genes (N=38). Survival analysis was performed using the Kaplan-Meier method and Cox proportional hazards regression with endpoint overall survival from time of biopsy to death of any cause.

**Results:** We found that the PCa-GI score had a bimodal distribution, identifying a distinct set of tumors with an activated GI expression pattern. Approximately 35% of samples were classified as PCa-GI high, which was concordant with prior reports. Liver metastases had the highest median score but after excluding liver samples, 29% of the remaining samples were still classified as PCa-GI high, suggesting a distinct phenotype not exclusive to liver metastases. No correlation was observed between GI score and proliferation, AR signaling, or NEPC scores. Furthermore, the PCa-GI score was not associated with genomic alterations in *AR, FOXA1, RB1, TP53* or *PTEN*. However, tumors with *MYC* amplifications showed significantly higher GI scores (p=0.0001). Patients with PCa-GI tumors had a shorter survival (HR=1.5 [1.1-2.1], p=0.02), but this result was not significant after adjusting for the liver as metastatic site (HR=1.2 [0.82-1.7], p=0.35). Patients with PCa-GI low samples had a better outcome after androgen receptor signaling inhibitors (ASI, abiraterone or enzalutamide) than other therapies (HR=0.37 [0.22-0.61], p=0.0001) while the benefit of ASI was smaller and non-significant for PCa-GI high samples (HR=0.55 [0.29-1.1], p=0.07). A differential pathway analysis identified FOXA2 signaling to be upregulated PCa-GI high tumors (FDR = 3.7 × 10^−13^).

**Conclusions:** The PCa-GI phenotype is prevalent in clinical mCRPC samples and may represent a distinct biological entity. PCa-GI tumors may respond less to ASI and could offer a strategy to study novel therapeutic targets.

## Introduction

Prostate cancer is a heterogeneous disease that ranges from indolent tumors without any need for therapy to rapidly progressing and lethal cancers. While most prostate cancers can be treated effectively by local therapy such as surgery or radiotherapy, metastatic prostate cancer is not curable and eventually develops treatment resistance. Prostate cancer is mainly driven by androgen receptor (AR) signaling and as a result, AR-targeted therapies are the main systemic treatment for patients throughout all stages of the disease^1^. However, metastatic prostate cancers universally develop resistance to AR treatment and the mechanisms of this resistance are complex and varied. Several of these mechanisms revolve around genetic alterations (such as mutations, amplifications, and gene rearrangement) of the *AR* gene^2-7^. Some tumors, however, exhibit reprogramming of the transcriptional state to an AR-independent phenotype and acquiring features of neuroendocrine prostate cancer (NEPC), or other lineages^8^. This has led to subtyping and classification efforts e.g. along the AR and NEPC axes^9^ or basal/luminal classifications^10^ which have been shown to be prognostic. While prostate cancer lineage plasticity has been most studied in the context of NEPC, other forms of lineage plasticity have been suggested including basal, stem-cell, and Wnt signaling-driven phenotypes^11^. Furthermore, a switch to a more specific gastrointestinal (GI) transcriptional phenotype has been suggested^12^, and may be detectable from a liquid biopsy analyzing circulating cell-free DNA with 5-hydroxymethylcytosine sequencing^13^. The GI phenotype was correlated with *SPINK1* expression (a pancreatic secretory trypsin inhibitor) that protects the GI tract from protease degradation. A subset of prostate cancers exhibits an outlier expression pattern of *SPINK*, suggesting its role as a potential driver and drug target in the disease^14,15^. The PCa-GI phenotype was recently described to be associated with a worse outcome after AR-targeted therapy and may be targeted with BET-inhibitors^16^. This study aimed to define the prevalence of the PCa-GI phenotype in metastatic castration-resistant prostate cancer (mCRPC) as well as assess the impact on prognosis and treatment response to AR-targeted therapy by analyzing the PCa-GI phenotype in 634 mCRPC samples with RNA-seq from four independent clinical cohorts.

## Methods

### Data cohorts

We used data from four previously published and publicly available mCRPC cohorts with RNA-seq data available. The RNA-seq data were previously combined and batch-effect corrected, and DNA alteration calls and survival data were used as previously described^10,17^. In total, the cohort consisted of 634 mCRPC samples from the West Coast Dream Team (WCDT, N=162^6,18,19^), the East Coast Dream Team (ECDT, N=266^5,20^), the Fred Hutchinson Cancer Research Center (FHCRC, N=157^21^), and the Weill Cornell Medical Center (WCM, N=49^22^). The WCDT and ECDT had survival data annotated. Details of this study cohort and the initial processing of the RNA samples, including the batch-effect correction, have been published previously^10^.

### Calculation of a gastrointestinal transcriptional score

The identification of a prostate cancer gastrointestinal (GI) score was previously published with a list of 129 genes that correlated with the phenotype and expression of *SPINK1* and a shorter list of 38 core genes^12^. For this analysis, we used the core set of 38 genes. We calculated the sum of z-scores for the 38 genes on the log2 transformed (with an offset of 8000 to avoid negative values) rank normalized and batch-effect corrected data. We used the same method to calculate pathway scores for the Cancer Hallmark pathways^23^, the Wikipathways^24,25^, and prostate-specific gene lists from the literature^22,26-30^. Based on the distribution of scores we dichotomized the PCa-GI scores into high and low samples with a cutoff at 0 for grouped analyses.

### Statistical Analysis

For correlation analysis, we used Spearman’s correlation. The Wilcoxon rank sum test was used to assess differences between groups unless otherwise noted. To assess differences between the PCa-GI low and PCa-GI high samples, we conducted a differential analysis of pathways using a Wilcoxon rank-sum test of pathways scores between the PCa-GI-high and PCa-GI low samples and calculated the median difference in pathway scores. P-values for the differential pathway analysis were adjusted for multiple testing using the False Discovery Rate (FDR) described by Benjamini and Hochberg^31^. Overall survival data from the time of biopsy to death of any cause was available for the ECDT and WCDT cohorts, and we used the Kaplan-Meier method to visualize survival probabilities, and the Cox proportional hazards model was used to assess survival differences between groups. All analyses were conducted in R, version 4.3.2.

## Results

### The prostate cancer gastrointestinal transcriptional phenotype (PCa-GI) is present and prevalent in clinical mCRPC biopsies

We calculated the PCa-GI score as the sum of z-scores for the 38 core genes and found that the PCa-GI score exhibited a bimodal distribution pattern. Given that several of the core genes are also expressed in hepatocytes, we performed the analysis excluding liver biopsy samples and observed a similar pattern (Figure 1A). All metastatic biopsy sites had individual samples with high PCa-GI score with the highest median score in liver biopsy samples (Figure 1B). Using a cutoff of 0, we found that 35% of all samples and 29% of non-liver samples had a high PCa-GI score. Furthermore, we plotted the sum of z-scores for the Hallmark Bile Acid Metabolism gene set^23^ against the PCa-GI score. Our analysis revealed that liver samples typically exhibited high scores for both gene sets. In contrast, non-liver samples often had elevated PCa-GI scores, but typically did not show high Bile Acid Metabolism scores, suggesting that the PCa-GI score identifies tumors distinct from the pathways typically activated in liver biopsy samples (Figure 1C). Finally, we examined publicly available single-cell RNA-seq data^32^ and examined the expression of the three PCa-GI associated genes *SPINK1, HNF1A*, and *HNF4G*. We found that their expression was specific to cancer cells and present at high levels in non-liver metastatic samples (Supplemental figure 1). Taken together, the PCa-GI phenotype is present in mCRPC samples and in both hepatic and non-hepatic metastases.

**Figure 1.**
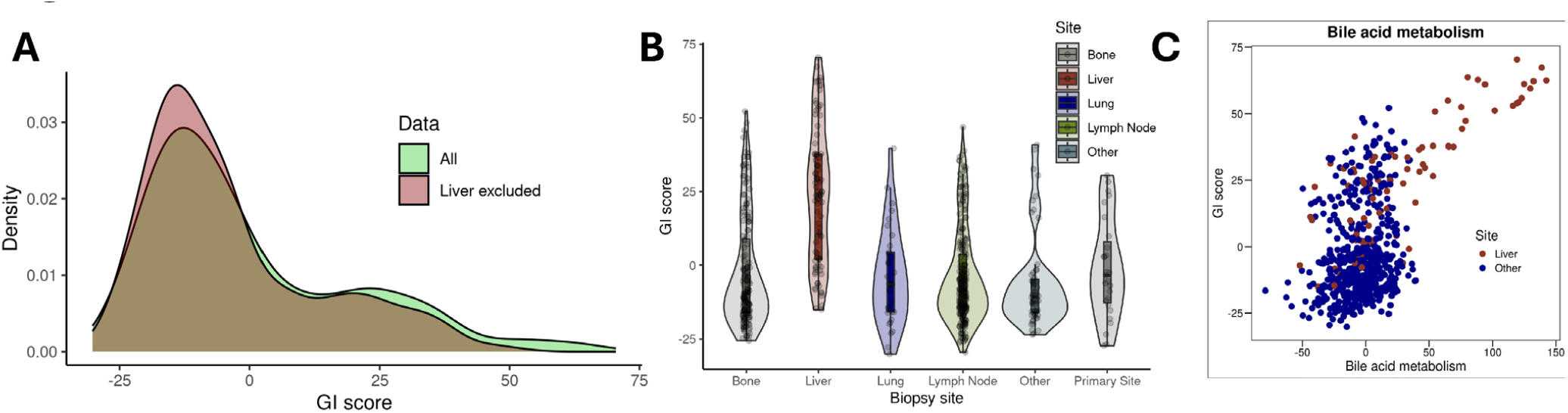
A gastrointestinal transcriptional phenotype exists among clinical mCRPC biopsy samples. A) Distribution of a transcriptional prostate cancer gastrointestinal (PCa-GI) score among 634 mCRPC biopsy samples in all (green) or non-liver (red) samples. B) GI scores in different metastatic biopsy sites. C) Scatterplot of GI score vs a bile acid metabolism score.

### The PCa-GI score is independent of established mCRPC drivers

Next, we sought to assess if the PCa-GI score was correlated with known drivers of aggressive mCRPC, such as AR-signaling, proliferation, and NEPC. We found no association with AR-signaling (rho = 0.02, p = 0.62), proliferation (rho = 0.024, p = 0.55) or NEPC (rho = 0.072, p = 0.07) (Figure 2A-C). There was also no association between the PCa-GI score and genomic alterations in *AR, FOXA1, PTEN, RB1*, or *TP53*, while there was a significantly higher PCa-GI score in *MYC* amplified tumors (p=0.000012) (Figure 2D). These results indicate that the PCa-GI is a distinct biological entity not captured by current transcriptional phenotyping approaches.

**Figure 2.**
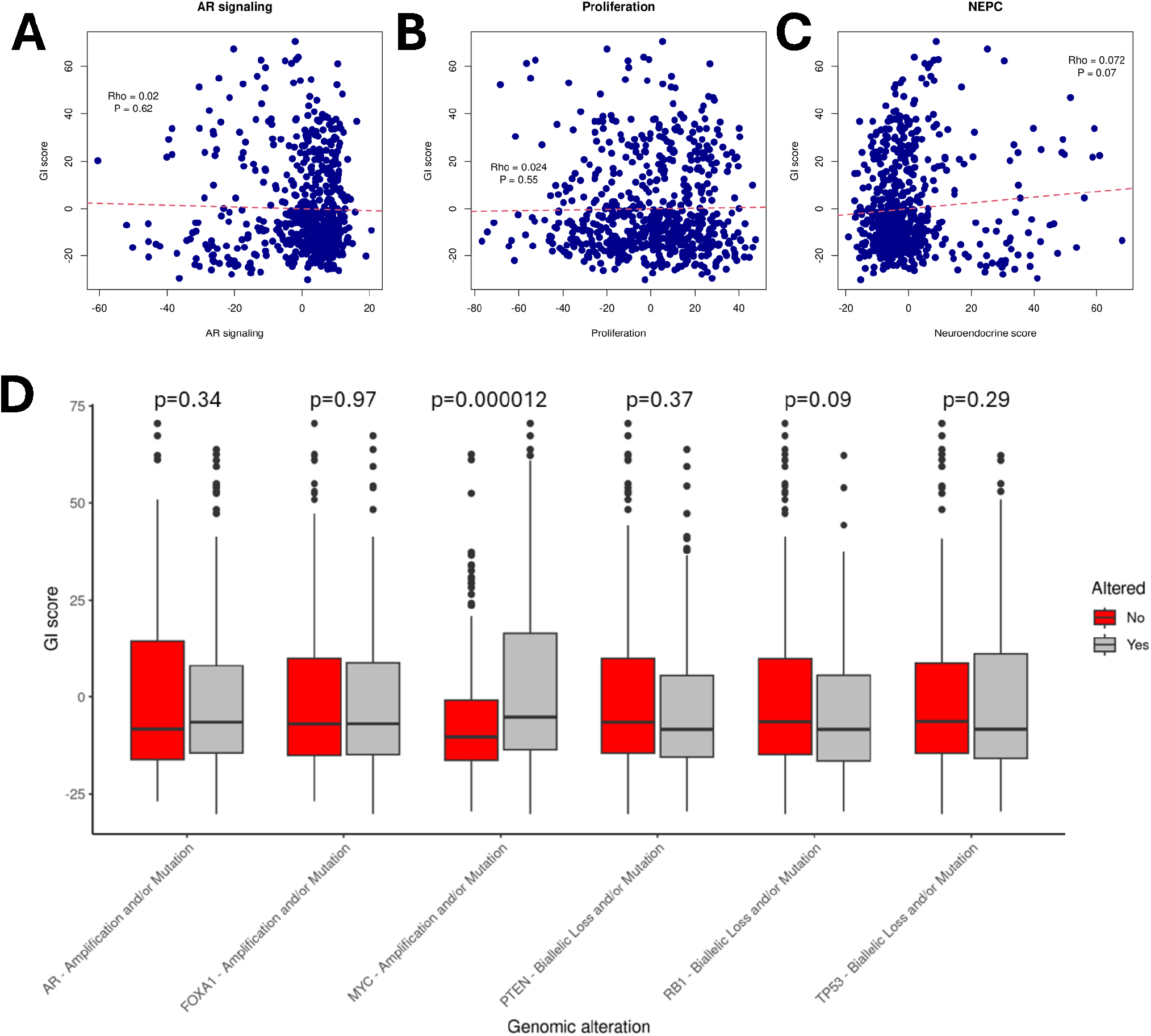
The prostate cancer gastrointestinal score represents a distinct biological entity. A) Scatterplot of the transcriptional gastrointestinal (GI) score and AR signaling. B) Scatterplot of the GI score and proliferation. C) Scatterplot of the GI score and neuroendocrine prostate cancer score (NEPC). D) The GI score and genomic alterations in *AR, FOXA1, MYC, PTEN, RB1*, and *TP53*.

### The impact of the PCa-GI phenotype on prognosis and survival after treatment with androgen signaling inhibitor (ASI)

We next evaluated the effect of the PCa-GI phenotype on survival in patients from the WCDT and ECDT cohorts, where the survival data was available. In the combined cohort the PCa-GI tumors had a worse survival than PCa-GI low tumors (HR = 1.5, 95%CI 1.1-2.1, p = 0.02) (Figure 3A). However, this effect was diminished after adjusting for the liver as a metastatic site (HR 1.2, 95%CI 0.82-1.7, p = 0.35). There was a numerically higher but not-significant PCa-GI score in tumors that had received a novel generation androgen signaling inhibitor (ASI) (p = 0.13) (Figure 3B). When stratifying the survival analysis by if the patients received immediate post-biopsy ASI, PCa-GI low tumors had better survival after ASI (HR = 0.37, 95%CI 0.22-0.61, p = 0.00011) (Figure 3C), while PCa-GI high tumors had a smaller and non-significant difference (HR = 0.55, 95%CI 0.29-1.1, p = 0.074) (Figure 3D). While recognizing that the numbers are low and the results hypothesis-generating, we did an exploratory analysis for the survival after post-biopsy ASI in ASI naïve tumors. In these ASI naïve tumors, PCa-GI low tumors had better survival after ASI than other subsequent therapies (HR = 0.2, 95% CI 0.073-0.54, p = 0.0016) while PCa-GI had no difference between ASI and other therapies (HR = 1.1, 95%CI 0.13-8.7, p = 0.95) (Supplementary Figure 2). A test for interaction between the PCa-GI phenotype and post-biopsy ASI among ASI naïve tumors showed statistical significance (p for interaction = 0.049), indicating that the PCa-GI status may predict clinical benefit of ASI therapy.

**Figure 3.**
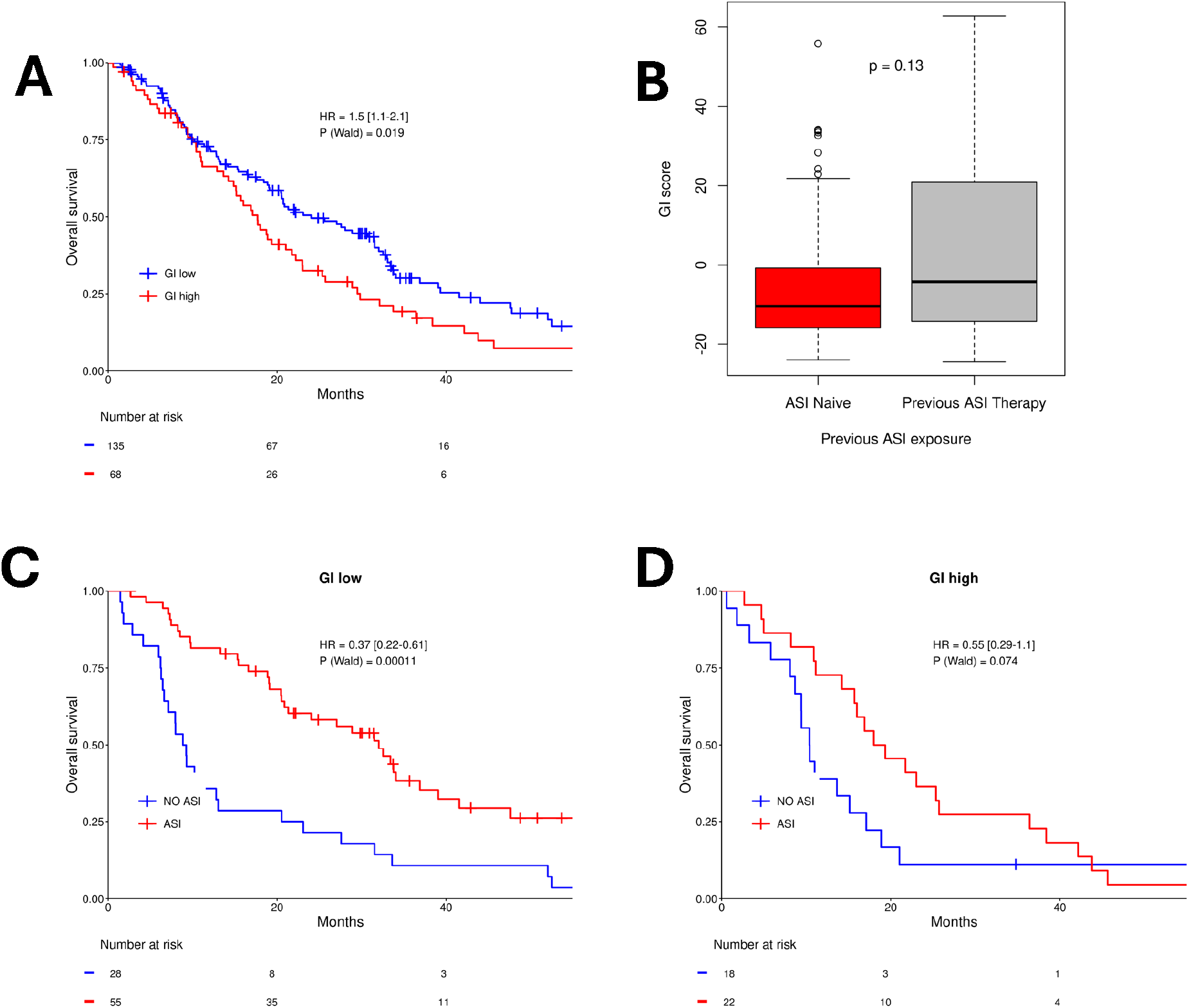
The prognostic and treatment predictive potential of the prostate cancer gastrointestinal score. A) Overall survival from time of biopsy for PCa-GI low and PCa-GI high tumors. B) Boxplot of PCa-GI score in tumors without or with prior androgen signaling inhibitor (ASI). C) Overall survival from time of biopsy depending on immediate postbiopsy ASI treatment or another postbiopsy therapy in PCa-GI low tumors. D) Overall survival from time of biopsy depending on immediate postbiopsy ASI treatment or another postbiopsy therapy in PCaGI high tumors.

### Potential biological drivers of the PCa-GI phenotype

Finally, we sought to explore biological differences and potential drivers of the PCa-GI phenotype. First, we plotted the correlation between the PCa-GI score and the Cancer Hallmark pathways, and strikingly the PCa-GI score did not correlate strongly with the Hallmark pathways, again suggesting that the PCa-GI phenotype is a distinct biological entity (Figure 4A). We next performed a differential pathway analysis between the PCa-GI high and PCa-GI low tumors (Figure 4B, full list of results of all tested pathways in Supplemental Table 1). As expected, the full PCa-GI score (using all 129 genes^12^) and the shorter core PCa-GI score (using the core 38 genes) were the most significantly different pathways. In addition, several liver-associated signatures that share common genes with the PCa-GI phenotype ranked high in the list. However, after these signatures, the top differential pathway was the FOXA2 pathway (Figure 4B). Interestingly, FOXA2 has recently been suggested as a driver of prostate cancer dedifferentiation and lineage plasticity^33^. In conclusion, these results suggest that the PCa-GI phenotype represents a distinct biological entity and that further studies may elucidate drivers and potential targets in this prostate cancer subtype.

**Figure 4.**
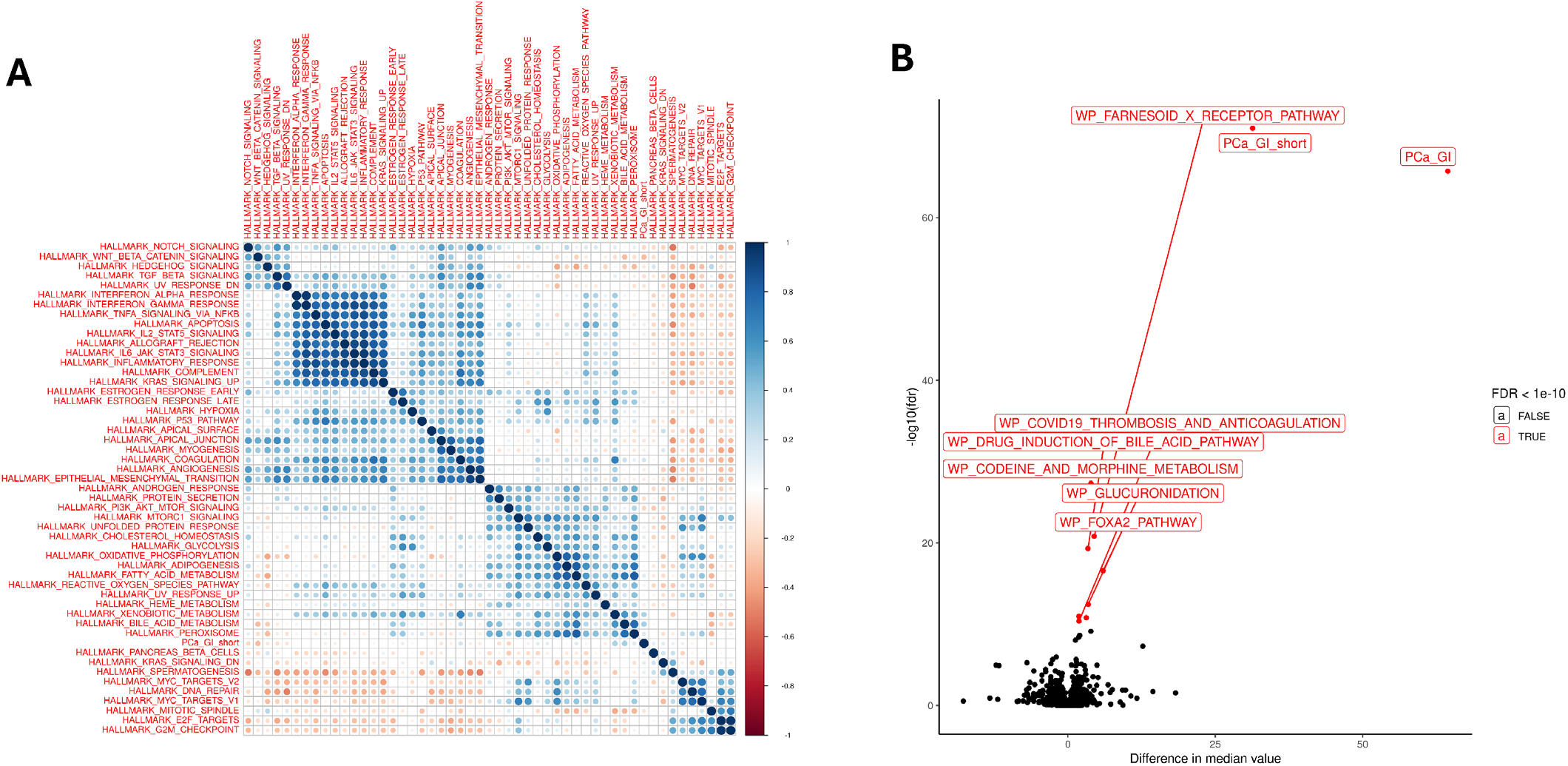
Biological differences in GI low and high tumors. A) Correlation plot of biological pathways in mCRPC. Each pathways score is calculated as the mean of Z-scores for genes in the pathways and the color in the plot depicts degree of inter pathway correlation. B) Differential pathway analysis between GI high and GI low tumors, excluding liver samples. The Y-axis represents the statistical strength as the -log10(false discovery rate) from a Wilcoxon rank sum test. The X-axis represents the difference in median pathway score.

## Discussion

Herein, we studied a previously described prostate cancer gastrointestinal (PCa-GI) phenotype in a large cohort of 634 clinical mCRPC biopsy samples. We showed that approximately 30% of mCRPC samples have the PCa-GI phenotype, which is in line with prior studies^12^. Importantly, while liver metastases exhibited the highest median PCa-GI scores, non-liver metastases can also express a PCa-GI transcriptional phenotype, and analysis of single-cell data demonstrated that PCa-GI related genes, such as *SPINK1, HNF1A*, and *HNF4G* were expressed in tumor cells in non-liver metastatic sites. The PCa-GI phenotype has been suggested to emerge under AR-targeted therapy and be resistant to further AR-targeted agents from *in vitro studies*^*12*^. Although the number of patients in this cohort with therapy information and outcomes data is small, the results herein support a potential association between the PCa-GI and resistance to ASI therapy. Finally, we showed that FOXA2 signaling is upregulated in PCa-GI high tumors demonstrating that further studies should be aimed at understanding the biological drivers of this phenotype for therapeutic exploitation.

The lack of correlation between the GI score and proliferation, AR signaling, or NEPC score suggests that PCa-GI phenotype in mCRPC may represent a distinct biological entity independent of the established prostate cancer progression markers. The absence of associations with genomic alterations in *AR, FOXA1, RB1, TP53*, or *PTEN* further suggests a different resistance mechanism than alterations in the *AR* gene and the development of aggressive variants or NEPC.

When analyzing survival outcomes with or without the PCa-GI phenotype, we only found a difference in survival with PCa-GI high tumors that was associated with the high prevalence of the PCa-GI in liver samples. More interestingly, while we did not observe a survival benefit in the overall population based on stratification of GI high and low patients, we did observe a statistically significant difference in survival after ASI treatment based on GI score in ASI naïve tumors. While the number of samples for this analysis is small, it mirrors preclinical data showing that activation of the PCa-GI phenotype by itself, does not alter tumor aggressiveness but makes it less responsive to AR-targeted therapy^12^. Together, these results nominate the PCa-GI phenotype as a potential target for developing novel therapies. Indeed, a recent study suggested that the PCa-GI phenotype may be targeted with BET-inhibitors^16^, and we are eagerly looking forward to future studies.

Although we analyzed a large cohort of clinical mCRPC samples, our study has limitations. Firstly, this study was based on an analysis of gene expression from RNA-sequencing, and while prior studies have confirmed the expression of PCa-GI related genes also at the protein level, the morphological and protein expression patterns should be determined in future studies. Secondly, this study was based on bulk RNA-sequencing, and the potential influence by non-tumor cells is difficult to analyze. However, we did observe the PCa-GI phenotype at all metastatic sites, and public single-cell data support the expression of PCa-GI related genes in tumor cells at different metastatic sites. As single-cell and spatial transcriptomics become more readily available, this should be further explored. Thirdly, these cohorts were not randomized and the analysis of survival after different therapies should be interpreted with caution. However, the results presented herein show the same trend as prior studies warranting further exploration of the PCa-GI phenotype as a mechanism of treatment resistance in mCPRC. Future studies should aim to delineate the molecular underpinnings of the FOXA2 pathway’s role in the GI phenotype and explore the therapeutic potential of targeting this pathway. Additionally, the mechanistic link between *MYC* amplification and the GI phenotype merits further exploration, potentially unraveling new avenues for therapeutic intervention.

## Conclusions

In conclusion, our study highlights that the PCa-GI phenotype is not related to androgen receptor signaling or neuroendocrine prostate cancer and may respond less to androgen receptor-targeted therapy. Further studies should be directed at elucidating and exploiting specific pathway alterations in the PCa-GI phenotype, potentially offering new insights into its oncogenic landscape and therapeutic vulnerabilities.

## Acknowledgements

We would like to acknowledge funding from the National Institutes of Health [grant numbers DP2 OD030734 to SGZ, 1UH2CA260389 to SGZ], Department of Defense [grant numbers PC190039 to SGZ, PC200334 to SGZ, HT94252310164 to MNS], Prostate Cancer Foundation (2022 Janssen—PCF Special Challenge Award to DAQ, 2022 Point Biopharma Young VAlor Investigator Award to MNS, 2021 Michael and Patricia Berns-PCF Young Investigator Award to MS), the Doris Duke Charitable Foundation (Physician Scientist Fellowship #2021088 to MNS), the Swedish Cancer Society (Cancerfonden, Junior Clinical Investigator Award to MS), the Swedish Prostate Cancer Foundation (Prostatacancerförbundet, to MS), and Hjelms stiftelse för medicinsk forskning (MS co-applicant).

## Supplementary figures

**Supplementary figure 1.**
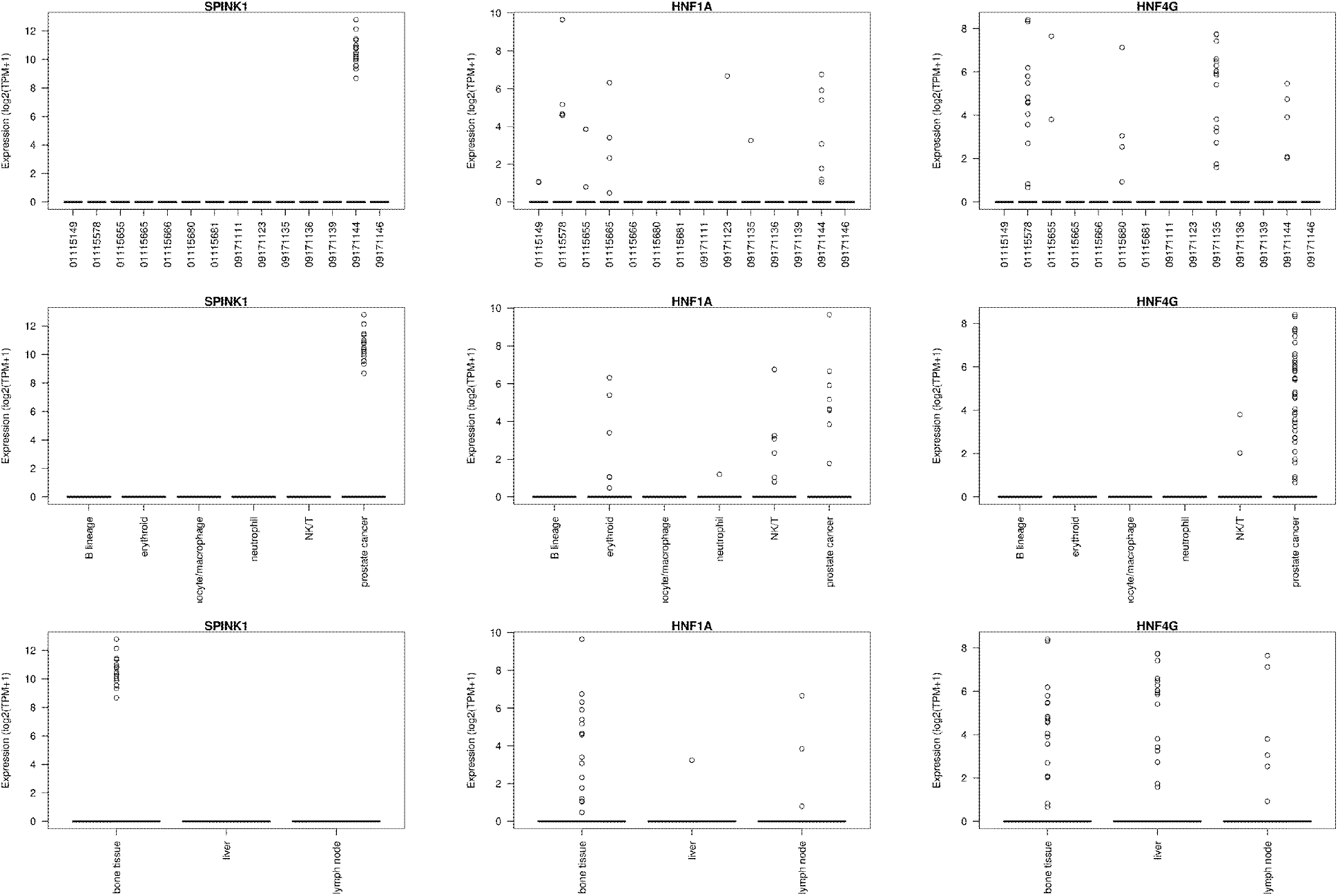
Expression of the gastrointestinal score associated genes *SPINK1, HNF1A*, and *HNF4G* in single-cell RNA-sequencing data from He et al.^32^. Top row is expression in single cells split per patient. Middle row is expression in single cells split per cell type. Bottom row is expression split per site of metastasis.

**Supplementary figure 2.**
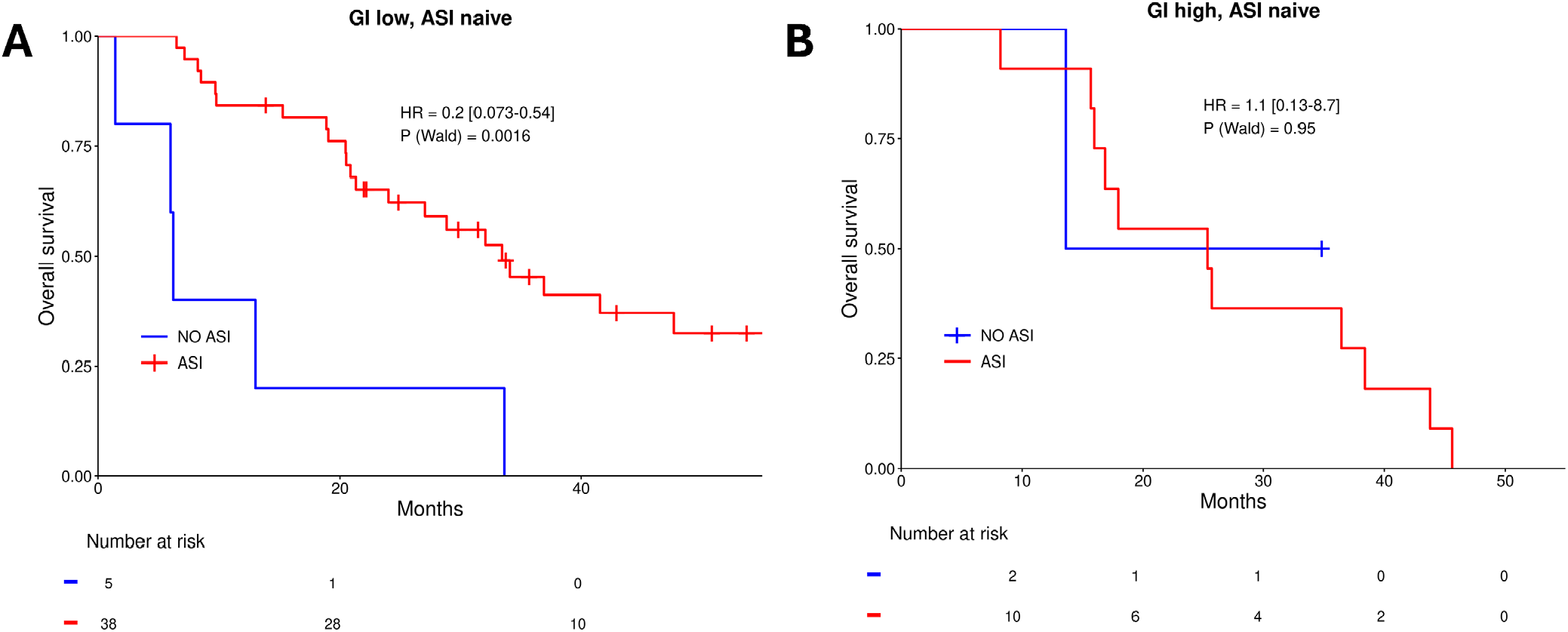
The treatment predictive potential of the prostate cancer gastrointestinal score in ASI naïve tumors. A) Overall survival from time of biopsy depending on immediate postbiopsy ASI treatment or another postbiopsy therapy in PCa-GI low and ASI naïve tumors. B) Overall survival from time of biopsy depending on immediate postbiopsy ASI treatment or another postbiopsy therapy in PCa-GI high and ASI naïve tumors.

## Supplementary tables

Supplementary table 1. Differential pathway analysis between GI low and GI high tumors. Cancer Hallmark, Wikipathways and additional prostate cancer relevant gene list from the literature were analyzed.

